# Tissue Specific Age Dependence of the Cell Receptors Involved in the SARS-CoV-2 Infection

**DOI:** 10.1101/2021.07.13.452256

**Authors:** Christian V. Forst, Lu Zeng, Qian Wang, Xianxiao Zhou, Sezen Vatansever, Zhidong Tu, Bin Zhang

## Abstract

The coronavirus disease 2019 (COVID-19) pandemic has affected tens of millions of individuals and caused hundreds of thousands of deaths worldwide. Due to its rapid surge, there is a shortage of information on viral behavior and host response after SARS-CoV-2 infection. Here we present a comprehensive, multiscale network analysis of the transcriptional response to the virus. We particularly focus on key-regulators, cell-receptors, and host-processes that are hijacked by the virus for its advantage. *ACE2*-controlled processes involve a key-regulator *CD300e* (a *TYROBP* receptor) and the activation of IL-2 pro-inflammatory cytokine signaling. We further investigate the age-dependency of such receptors and identify the adipose and the brain as potentially contributing tissues for the disease’s severity in old patients. In contrast, several other tissues in the young population are more susceptible to SARS-CoV-2 infection. In summary, this present study provides novel insights into the gene regulatory organization during the SARS-CoV-2 infection and the tissue-specific age dependence of the cell receptors involved in COVID-19.

## Introduction

On December 31, 2019, the WHO was notified about a cluster of novel pneumonia cases in Wuhan City, Hubei Province of China. The causative agent was linked to a novel by Chinese authorities on January 7, 2020, inducing the activation of the R&D Blueprint as part of WHO’s response to the outbreak. Coronaviruses (CoVs) belong to the group of enveloped, single, positive-stranded RNA viruses causing mild to severe respiratory illnesses in humans^1^. In the past two decades, two worldwide outbreaks have originated from CoVs (SARS, MERS) capable of infecting the lower respiratory tract, resulting in heightened pathogenicity and high mortality rates^2^. We are currently amid a third pandemic caused by a new CoV strain, the severe acute respiratory syndrome coronavirus 2 (SARS-CoV-2), the causative agent of coronavirus disease 2019 (COVID-19). In the majority of cases, patients exhibit either no or mild symptoms, whereas in more severe cases, patients may develop severe lung injury and die from respiratory failure^2,3^.

A viral infection generally triggers a physiological response at the cellular level after the initial replication of the virus^4^. The cellular system has an arsenal of pattern recognition receptors (PRRs)^5^ at its deposal that guard against various microbes inside and outside of the cell. PRRs bind distinct structural features that are conserved among different pathogens^6^. In a viral infection, intracellular PRRs are detecting viral RNA defective particles that are often formed during virus replication^7^. Pathogen detection assembles the initial steps of a signaling cascade to activate downstream transcription factors, such as interferon regulator factors (IRFs) and nuclear factor kB (NF-κB)^6,8^, which causes the activation of two general antiviral processes^6^. The first, predominantly intracellular, process initiates cellular defenses via transcriptional induction of type I and III interferons (IFN-I and IFN-III, respectively). Subsequently, IFN upregulates IFN-stimulated genes (ISGs) with antiviral properties^9^. The second, inter-cellular cascade of antiviral counteraction refers to the recruitment and coordination of a multitude of leukocytes. Chemokine secretion^10,11^ orchestrates this concerted action of immune-system countermeasures. The selection pressure induced by such a broad antiviral response of the host and the evolvability of viruses has resulted in countless viral countermeasures^12^. Thus, the host response to a virus is generally not uniform. Viral infections can cause a spectrum of various degrees of morbidity and mortality.

Indeed, additional factors, such as sex, age, other genetic factors, contribute to the diversity of immune response. Concerning COVID-19, age has been identified as the most significant risk factor in the mortality of patients. The overall symptomatic case fatality risk (the probability of dying after developing symptoms) of COVID-19 in Wuhan was 1.4% (0.9–2.1%) as of February 29, 2020. Compared to those aged 30–59 years, those aged below 30 and above 59 years were 0.6 (0.3–1.1) and 5.1 (4.2–6.1) times more likely to die after developing symptoms^13^. Similar data were reported for the United States. From February 12 to March 16, 2020, the Center for Disease Control (CDC) estimated a case-fatality rate of patients 55-64 years old with 1.4 – 2%. This rate was 10.4 – 27.3% for patients 85 years or older^14^.

To better understand the disease’s molecular basis, we sought to characterize the transcriptional response to infection in both in vitro cell systems (tissue cultures and primary cells) and *in vivo* samples derived from COVID-19 patients. We employed an integrative network-based approach to identify host response co-expression networks in SARS-CoV-2 infection. In particular, we investigated functional processes and key regulators affected by this specific virus, receptors used for entry, and processes hijacked for enabling viral life cycles. We further studied the age-dependence of targets, mainly receptors that the virus utilizes for entry and its life cycle.

## Results

RNA-seq data from cell lines (NHBE, Normal Human Bronchial Epithelial cells, A549, adenocarcinomic human alveolar basal epithelial cells, and Calu-3, lung adenocarcinoma epithelial cells) and lung biopsies of two patients infected by SARS-CoV-2 were recently made available on NCBI/GEO (GSE147507)^6^. A second, clinical, transcriptomic dataset for a cohort of COVID-19 patients together with uninfected controls has recently been published^15^. Data were obtained from bronchoalveolar lavage fluid (BALF) and PBMCs (10 samples total: 3 PBMC control, 2×2 BALF infected, 3 PBMC infected). RNA-seq data is available through the Beijing Institute of Genomics (BIG) Data Center (https://bigd.big.ac.cn/) under the accession number: CRA002390. We have combined the BALF with the lung biopsy datasets after batch correction, yielding datasets containing a total of 11 samples (6 infected and five control). These datasets were processed by an integrative network analysis approach. Data from PBMCs and cell lines were excluded. For validation purposes, we have further secured data from a second cohort of 142 patients from the NYU Langone Health Manhattan campus that required invasive mechanical ventilation^16^.

### Integrative network biology analysis of the β-coronavirus – host system

The basis of our prediction of SARS-CoV-2 processes and the host response is an integrative network analysis approach that combines network inference and network topological methods with molecular signatures. We first identified differentially expressed genes (DEGs) in each dataset that showed significant changes during SARS-CoV-2 infection. The biological functions of DEG signatures from each dataset were assessed by gene-set enrichment methods. Given the particular interest of human patients’ COVID-19 response, we used a corresponding subset of transcriptome data to infer multiscale gene co-expression MEGENA networks. We ranked MEGENA network modules based on their enrichment for DEGs. MEGENA modules were functionally assessed by GO, MSigDB, and blood cell-type-specific gene-sets. We also investigated the underlying network topological structure by testing the network neighborhood of target genes for enrichment by SARS-CoV-2 DEGs and signatures responding to ACE2 overexpression. Finally, we analyzed the age-dependency of molecular processes during SARS-CoV-2 infection by employing a linear regression model on baseline gene expression using Genotype-Tissue Expression (GTEx) data.

### Molecular signatures of SARS-CoV-2 infection

We have identified 572 up-, and 1338 downregulated DEGs from patient-derived lung biopsy, as well as 3,573 up- and 1,630 downregulated DEGs from human patient BALF expression data. 2,382 DEGs are upregulated, and 2,526 DEGs are downregulated in A549 cell lines (2,017 up- and 2,354 downregulated in Calu3 cell lines, resp.). The exceptions are the NHBE and the first batch (Series 2) of the A549 data (GSE147507), which yielded a fraction of significant DEGs, with 144 genes up- and 55 genes downregulated in NHBE cells as well as 88 genes up- and 14 genes downregulated in A549 (Series 2). All datasets have comparable numbers of samples. DEGs were considered significant with FDR ≤ 0.05 and a fold change of 1.5 or higher.

As others have already noted^6^, there is a lack of *ACE2* expression in cell line data. A key-protein relevant for SARS-CoV-2 entry as well as an ISG, *ACE2* is not significantly expressed in cell lines (S5_A549: 3.2 fold, FDR = 0.15; Calu3: 0.77 fold, FDR = 0.12; NHBE: 1.2 fold, FDR = 0.52). Only in the lung biopsy (27.6 fold, FDR = 3.70 e-06) and in BALF (50.5 fold, FDR = 0.066), we were able to identify significant expression fold change between healthy/Mock control and infection. According to GTEx data, ACE2 baseline expression is observed in the small intestine (Terminal Ileum), female breast, thyroid, subcutaneous adipose tissue, testis, and coronary artery (**Table S1**). A detailed, single-cell-based study identified that *ACE2* and *TMPRSS2* are primarily expressed in bronchial transient secretory cells^17^. *TMPRSS2* expression is inconsistent in our datasets. It is highly upregulated in BALF (47.2 fold, FDR = 2.98 e-04) and upregulated in Calu3 cells (2.13 fold, FDR = 2.71 e-03), but downregulated in lung biopsy samples (0.16 fold, FDR = 8.91 e-07). As we are mostly interested in an organismal response, our primary focus is on samples of human patients.

To validate our findings, we compared DEGs called during our analysis of human patients samples and results from the NYU COVID-19 study^16^. For this purpose, we employed Super Exact Test^18^, a generalization of Fisher’s Exact Test to evaluate the set-overlap of multiple sets. BALF and lung biopsy data show significant overlap with NYU COVID-19 data (Figure S1).

### Receptors, host-factors and biological processes required for the viral life cycle

Given that *ACE2* is essential for SARS-CoV-2 entry^19^, and further, the viral life cycle, we hypothesize that *ACE2* expression may trigger other processes relevant to the viral life cycle. As we have established in the previous section that *ACE2* is indeed upregulated in human lung samples (both BALF and lung biopsy), we were interested in the effect of ACE2 expression. To determine which receptors and targets are involved in such processes, we performed a network enrichment analysis using the ACE2 overexpression signatures from the Blanco-Melo *et al*. dataset^6^ and identified genes that potentially serve as novel host receptors and targets facilitating the entry of the SARS-CoV-2 into the host cell. For this purpose, we constructed a multiscale co-expression network to investigate co-expression and co-regulation relationships among genes underlying SARS-CoV-2 infection. In particular, we were interested in the organismal response from patients infected by SARS-CoV-2. Thus, we combined the available datasets from BALF and lung biopsies to construct a multiscale co-expression network of 13,398 genes and 35,483 interactions using MEGENA^20^ (**Figure 1A**). This co-expression network includes 900 modules. The majority of the top-ranked modules (using DEGs from both patient and cell data by excluding the ACE2 overexpression (ACE2oe) dataset; see methods section) are enriched for well-known biological functions related to viral infection, including cell cycle, ribosome/translation, NF-κB canonical pathway, or cytokine signaling. The 20 top-ranked modules are shown in **Figure 1B** as a sector of a circus plot, together with information on enrichment for up and downregulated DEGs and signature sets (MSigDB, blood cells, ARCHS^4^ tissues, and cell lines, SARS-CoV-2 life cycle genes, inflammasome, ISGs, transcription factors, miRNA targets). A few of these modules are enriched for MSigDB functions (**Figure 1C**). As expected, we have identified a variety of cell types from the ARCHS^4^ database accordant to the infection scenario, ranging from lung tissue and epithelial cells (**Figure 1D**), aveolar macrophages as well as lymphocytes (**Figure 1D**).The enrichment for the two main DEG signature sets, BALF and human lung biopsy are shown in **Figures 1E** and **1F**. Although there are differences in the DEGs between these two DEG sets, we have identified common DEG enrichment in modules M2, M9, M12, M66, M68 and M400. Most of these modules are related to translation and the ribosome.

**Figure 1.**
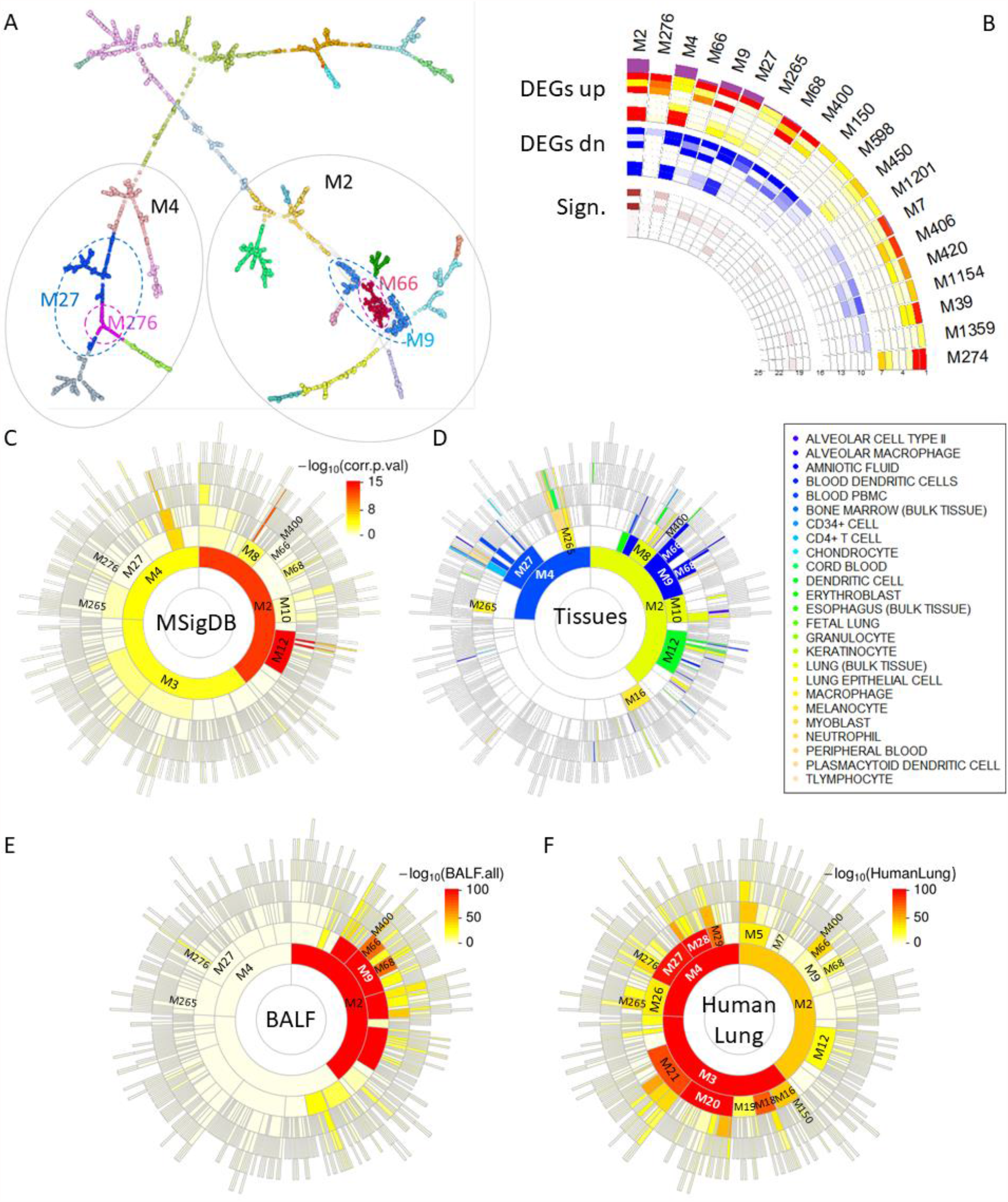
Gene co-expression modules associated with SARS-CoV-2 infection. (A) A global MEGENA network. Different colors represent the modules at one particular compactness scale. (B) The top 20 MEGENA modules most enriched for the SARS-CoV-2 up- and downregulated DEG signatures are shown (outer rings: “DEGs up” and “DEGs dn,” resp.). The center rings (“Sign.”) show additional signatures, including biological processes, cells, and tissues, as well as SARS-CoV-2 host factors based on PPI. (C) A Sunburst plot of all 934 modules enriched for MSigDB canonical processes (C2.CP) is shown. (D) The module enrichment for 25 lung pathology-related tissue signatures after the “ARCHS^4^ ” database^52^ is depicted. (E, F) Sunburst plots of module enrichment for DEGs concerning (E) BALF and (F) lung biopsy tissues are displayed. The color bars in (C, E, and F) show the negative decadic logarithm of the adjusted P-values.

**Figure 2A** shows a heat map of the 30 best-ranked receptors, along with fold change (FC) of expression during SARS-CoV-2 infection in lung samples and cell lines. All the targets are members of the M2-M10-M77 branch, except for *BTK* and *THEMIS2* (M2-M8-M59 branch) and *EXOC7* and *PTPRM* (M3-M20-M203 branch). Module M10, together with ACE2oe signature genes, is shown in **Figure 2B** (**Figure S2** depicts parent module M2). As shown in **Figure 1B**, M2, M10 and M77 are highly enriched for the ACE2oe signature with FET P-value = 1.20e-95 (1.7 Fold enrichment (FE)), 1.54e-20 (2.1FE) and 7.88e-13 (2.7 FE). All three modules are further enriched for lung tissue signatures after ARCHS^4^ tissues. Other modules such as M4, M9, M66, M69, M265, and M450 are also significantly enriched for ACE2oe signature (**Figure 2C**). M2 (rank 1) and M4 (rank 3) are the two largest modules associated with SARS-CoV-2 infection. They are associated with different biological functions such as ribosome (M2) and transcription (M4) (**Table S2**). M2 and M4 are the parents of several daughter modules. For example, in addition to the modules mentioned above, M10 (rank 35. **Figure 3A**) and M77 (rank 38, **Figure 3B**), M2’s daughter modules include highly ranked M7 (rank 14), M9 (rank 5, **Figure 3C**), M66 (rank 4, **Figure 3D**), M68 (rank 8), M400 (rank 9), M450 (rank 12), and M1201 (rank 13). A few of these modules are enriched for MSigDB functions (**Figure 1C**). Module M7 is enriched for phenylalanine metabolism, M9 for epithelium development and IL-2 signaling, M10 developmental biology, M68 meiotic recombination, and nucleosome assembly. Although M66, M400, M450, and M1201 are best-ranked and enriched for SARS-CoV-2 signatures, they are not significantly enriched for any known biological functions. Thus, these modules potentially indicate novel biological processes relevant to COVID-19. For example, the fourth-ranked M66 is driven by downregulated key regulators *DOHH, TMEM201* (or *SAMP1*), *TNFRSF25*, and *ZNF419*, as well as upregulated *ENTPD3* and *IFITM1* (**Figure 3D**).*TMEM201* is required for mitotic spindle assembly and γ-tubulin localization. The depletion of *TMEM201* results in aneuploidy phenotypes, i.e., the presence of an abnormal number of chromosomes in a cell, yielding bi-nucleated cells, and failed cytokinesis^21^. *TNFRSF25* is a member of the TNF-receptor family. This receptor has been shown to stimulate NF-κB activity and regulate cell apoptosis. *TNFRSF25* is further thought to be involved in controlling lymphocyte proliferation induced by T-cell activation. Thus, M66 likely plays a role in cytokinesis and cell proliferation. Concerning M4, highly ranked sub-modules (children) are M27 (rank 6, **Figure 3E**), M265 (rank 7), M276 (rank 2, **Figure 3F**). M276, with 81 genes, includes upregulated hemoglobin subunits δ, γ1, and μ (*HBD, HBG1, HBM*), which form part of the hemoglobin complex (FET P-value = 0.05, 62.1 FE). M276 is potentially responsible for oxygen transport (FET P-value = 0.089, 49.7 FE). M27 and M265 are not significantly enriched for any biological function (**Figure 1A** shows the M4-M27-M276 branch).

**Figure 2.**
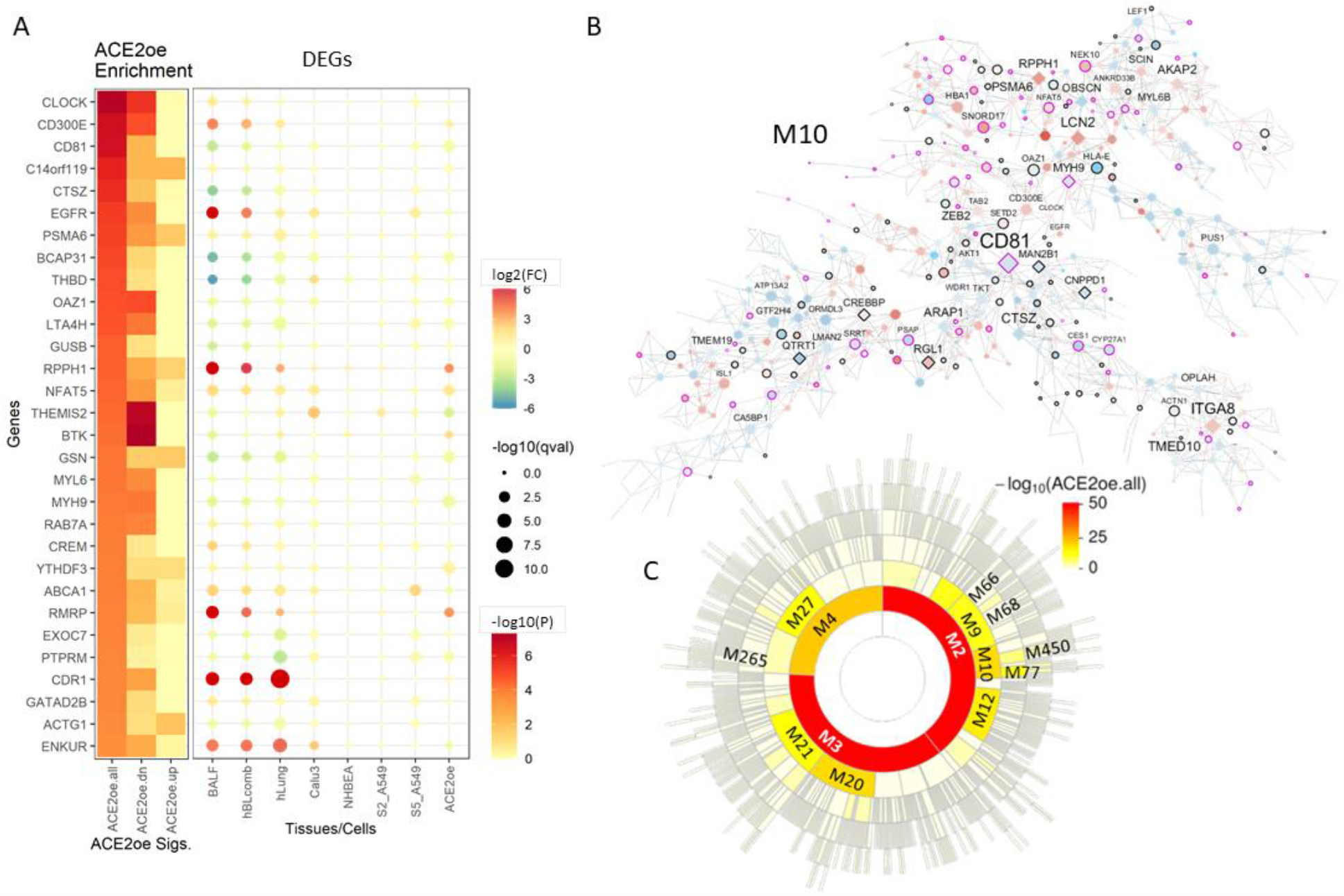
Network neighborhood and network enrichment for gene signatures and key regulators. (A) Top-scored targets after network enrichment by ACE2 overexpression signatures together with their directional response are shown. Many of these targets are members of M10 (B). The color tiles refer to network enrichment scores. The “-log10(P)” color scale on the right refers to the cumulative P-value used for ranking. Dark red color denotes a higher rank. The bubble plot denotes up- (red) and downregulated (blue) genes. The color of the circles refers to the fold change of expression between virus-infected and mock-infected samples. The size indicates the FDR as –log10(qval). (C) The number 35 ranked module M10 is depicted, which is significantly enriched for ACE2oe signatures. The node color indicates a directional response. Red nodes are upregulated, blue nodes are downregulated after infection. Diamond-shaped nodes indicate key regulators. The nodes with a black border denote genes significantly responding to ACE2 overexpression with fold change (FC) of 1.5 or higher. Purple borders indicate ACE2oe responding genes with FC ≥ 2. (E) A sunburst plot of the modules with ACE2oe enrichment is shown.

**Figure 3.**
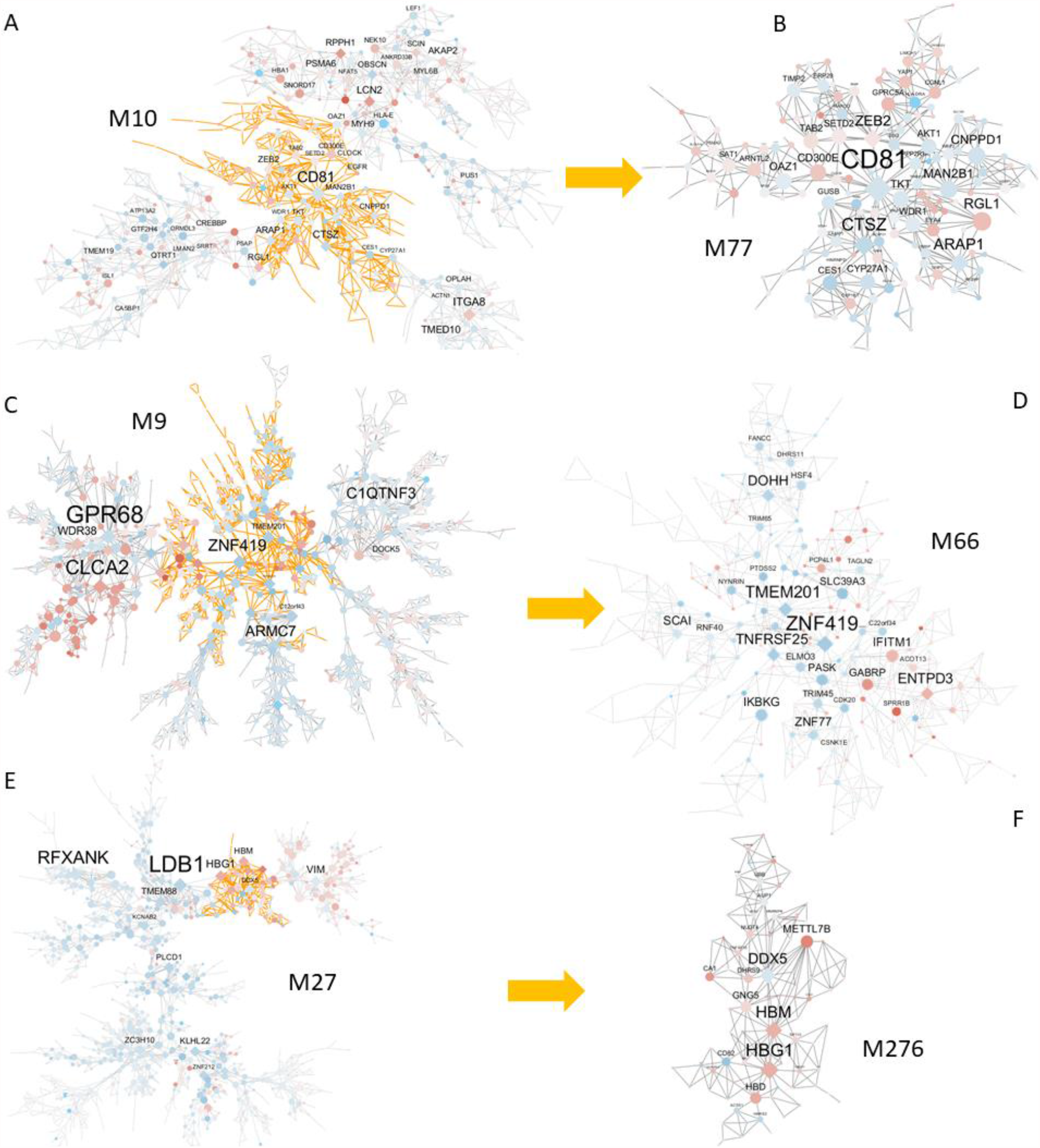
Gene co-expression modules associated with SARS-CoV-2 infection. (A) With rank 35, M10 is not among the best 20 ranked modules. It is potentially responsible for cellular stress response/Golgi apparatus/antigen processing and presentation and is enriched for DEGs, ACE2oe, and bulk lung tissue signatures. (B) Number 38 ranked module M77 is a daughter module of M10. M77 potentially functions for the regulation of cell adhesion. Like its parent module M10, M777 is enriched for DEGs, ACE2oe, and bulk lung tissue signatures. (C) M9 is the parent of M66 and ranked number 5, and is enriched for DEGs and ACE2oe signatures. Similar to M66, it is enriched for macrophages/neutrophils tissue signature. (D) Ranked fourth and second-ranked module with less than 100 genes is M66, which is enriched for DEGs and ACE2oe signatures. M66 is enriched for macrophages/neutrophils ARCHS^4^ signature. (E) M27 is the parent of M276 and ranked sixth. It is enriched for DEGs, ACE2oe, and blood PBMC signatures. (F) The top-ranked module with less than 1000 genes, M276, is highly enriched for upregulated DEGs. M276 is among the smallest top-ranked modules with 81 genes. – Node colors refer to the direction of regulation. Upregulated genes are red, and downregulated genes are blue. Diamond-shaped nodes denote key regulators. The size of the nodes refers to the connectivity in the network. (A, C, E) The subnetworks with orange edges refer to the corresponding daughter modules shown in (B, D, F).

The best-ranked ACE2oe network enriched targets are *CLOCK, CD300e, CD81, C14orf119*, and *CTSZ*. All but *C14orf119* are in the immediate network neighborhood of *CD81* (see **Figure 3F**). Clock circadian regulator (*CLOCK*) plays a central role in the regulation of circadian rhythms. *CLOCK*, a transcription factor, is upregulated in BALF and A549 samples.*CD300e* is a member of the *CD300* glycoprotein family of transmembrane cell surface proteins expressed on myeloid cells. It is upregulated in lung samples. The protein interacts with the TYRO protein tyrosine kinase binding protein (*TYROBP*) and is thought to act as an activating receptor. Activation via *CD300e* provided survival signals that prevented monocyte and Myeloid dendritic cells apoptosis, triggered the production of pro-inflammatory cytokines, and upregulated the expression of cell surface co-stimulatory molecules in both cell types^22^. The expression and function of human *CD300* receptors on blood circulating mononuclear cells are distinct in neonates and adults^23^, potentially contributing to the difference in clinical outcome after COVID-19 infection. Zenarruzabeitia *et al*. reported a stark down-regulation of CD300e on monocytes in patients with severe disease. However, we cannot confirm this finding in our BALF validation data. In the NYU COVID-19 study, CD300e is upregulated 1.6 fold in patients with severe diseases compared to patients with a mild outcome. Another ACE2oe network enriched target is *CD81*, with down-regulation in lung samples and cell-lines. *CD81* is an entry co-receptor for the Hepatitis C virus. *CD81* is the only ACE2oe target which network neighborhood is significantly enriched for SARS-CoV-2 signatures, yielding a rank of 79 based on NWes. Furthermore, *CD81* is a key regulator in the M2-M10-M77 branch (**Figures S2, 3A, and 3B**). Thus, *CD81* is potentially a novel host cell receptor that SARS-CoV-2 requires for entry and, therefore, a therapeutic target. Cathepsin Z (*CTSZ*) is a lysosomal cysteine proteinase and member of the peptidase C1 family. It is downregulated in lung samples and slightly upregulated in A549. Similar to *CD81, CTSZ* is a key regulator in M2-M1-M77. Singh et al., 2020 hypothesized that cathepsins are among other factors facilitating SARS-CoV-2 entry into the host cell^24^. The epidermal growth receptor *EGFR* is a transmembrane glycoprotein and present on the cell surface of epithelial cells. It is significantly upregulated in lung samples, A549, and Calu3 cells. *EGFR* is a host factor for hepatitis C virus entry^25^. Respiratory viruses induce *EGFR* activation, suppressing IFN regulatory factor (IRF) 1–induced IFN-λ, and antiviral defense in airway epithelium^26^. Thus, *EGFR* may not be required for SARS-CoV-2 entry, but it may be a potential host factor for the viral life cycle.

We validated our findings with results derived from the NYU COVID-19 cohort. Figure S3A shows a heatmap of 20 best ranked modules enriched for DEG signatures identified in this manuscript and deduced from the NYU COVID-19 cohort. Although the majority of modules is enriched for the combined lung and BALF data set, we can identify significant enrichment for best-ranked modules, in particular, for NYU COVID-19 DL and HL signatures. We further evaluated the similarity in gene content between modules from this study and modules derived from the NYU COVID-19 cohort (Figure S3B). In particular, best-ranked modules show significant overlap, validating the findings.

We have further investigated other cell-surface proteins, in particular cell surface receptors. For this purpose, we use data on experimentally verified high-confidence cell surface receptors from the cell surface protein atlas^27^ and data from the in silico human surfaceome^28^ – an extension from the protein atlas by using the measured protein data as a learning set for in silico prediction. From 2800 surface proteins, 1199 are classified as receptors by Surfaceome^28^, capable of transducing signals triggered by binding ligands or, hypothetically, surface proteins of the SARS-CoV-2 virion. Similar to the behavior of *ACE2*, we hypothesize that the expression of genes coding for such surface proteins can be triggered by the infection. We further hypothesize that such surface proteins mediate the transcriptomic response of downstream genes. Thus we expect up-regulation of the surface protein-coding genes and enrichment of DEGs in such receptors’ network neighborhood. Out of the 1199 receptors from the Surfaceome, 413 are in the MEGENA network. We identified further candidates in addition to the above-discussed surface receptors and key regulators *CD81, CD300E*, and *EGFR*. We expanded our criteria and included surface proteins that are significantly expressed across all datasets (employing ACAT, an aggregated Cauchy association test^29^). Surface proteins with the lowest aggregated P-value that are upregulated in most datasets were chosen. The highest-ranked candidate is lysosome-associated membrane glycoprotein 3 (*LAMP3*), followed by *EGFR*, as discussed above. LAMPs family plays a critical role in the autolysosome fusion process. *LAMP3* is expressed explicitly in lung tissues and is involved in influenza A virus replication in A549 cells^30^. It activates the PI3K/AKT pathway required for the influenza life cycle and necessary for SARS-CoV to establish infection, as demonstrated in Vero E6 cells^31^. Third-best ranked surface protein is CEA cell adhesion molecule 1 (*CEACAM1*). Multiple cellular activities have been attributed to the encoded protein, including roles in the differentiation and arrangement of three-dimensional tissue structure, angiogenesis, apoptosis, tumor suppression, metastasis, and the modulation of innate and adaptive immune responses. Both *CEACAM1* and *LAMP3* are members of the M4-M27 branch.

### SARS-CoV-2 triggered surface protein receptors expression show clear tissue-specific age-dependency

We were also interested in the age-dependency of the molecular processes involved in SARS-CoV-2 infection. A significant age disparity for severe cases, often causing death, has been widely reported for COVID-19. Being highly disproportional, more elderly patients experience severe symptoms and die due to this particular disease. We hypothesize that many host factors required for the virus life cycle have an age-dependent expression. By filtering in the genes upregulated in at least two of the SARS-CoV-2 studies, we obtained 213 genes encoding cell-surface proteins. These surface proteins are involved in transmembrane transport of small molecules (MSigDB c2.cp enrichment: P = 3.38e-08, 4.3 fold), ERBB(4) network pathway (P = 7.46e-08, 5.5 fold), neuroactive ligand-receptor interaction (P = 8.51e-05, 4.7 fold) or cytokine-cytokine receptor interaction (P = 1.22e-04, 4.2 fold). The tissue-specific age-dependency of these genes’ baseline expression was calculated by a linear model using data from GTEx (see **Methods**). We examined correlations between the expression of these SARS-CoV-2 triggered surface protein receptors (STSPRs) with chronological age using GTEx v8 data covering 46 tissues (**Table S3**). A large number of these surface protein receptors have their gene expression levels associated with age in many tissues, especially in the tibial artery, tibial nerve, and visceral fat. More than 70 receptors were significantly correlated with age. In contrast, very few receptors were associated with age in the liver, coronary artery, and brain substantia nigra (<5 receptors). Moreover, in most cases, the gene expression levels of these receptors were increased with age (**Table S4**).

We further examined the overall correlation between STSPRSTSPRs expression and age in a tissue-specific manner. Specifically, we first computed a composite receptor score (CRS) for each tissue of each sample in GTEx by summarizing the normalized expression values of the STSPRSTSPRs and then assessed the correlation between CRS and age (see **Methods** for details; **Figure 4)**. Three tissues, including the tibial artery, skeletal muscle, and subcutaneous fat, show the strongest positive correlations between their respective CRS and age. On the other hand, the whole blood, the frontal cortex (BA9), the ovary, and the cerebellum have the strongest negative correlations. Interestingly, the lung is ranked 31 out of 46 tissues, indicating that COVID-19 may impact far more tissues in different age populations than what we observed. As expected, the top-ranked tissues have a large number of significantly age-correlated receptors, consistent with the direction of the overall correlation. For example, in the tibial artery, which has a significant positive CRS-age correlation, 94 STSPRSTSPRs are significantly positive, and nine STSPRSTSPRs are significantly negatively correlated with age. Whereas in the frontal cortex, 56 STSPRs are significantly negative, and two STSPRs are significantly positively correlated with age, respectively (**Figure 4A**). The age effect on various disease pathologies is known for some of these tissues, with significant correlations between CRS and age. For example, age is a known risk factor for adverse outcomes in peripheral artery disease. The risk of severe limb ischemia, the sudden loss of blood flow to a limb caused by embolism or thrombosis, significantly increases with age^32^. Thrombosis and microvascular injury have been identified as an implication of severe COVID-19 infection^33^. Another example is skeletal muscles with well-studied age-related wasting and weakness. Cellular and molecular mechanisms contributing to a decline in muscular function involve neuromuscular factors, hormones, testosterone or growth hormone, insulin, myogenic regulatory factors (MRFs), the Notch signaling pathway, as well as cytokines and inflammatory pathways^34^. A cytokine storm and robust production of cytokines^6^ are known to contribute to the severity of COVID-19 infections^35^, potentially inducing systemic effects across many tissues and organs.

**Figure 4.**
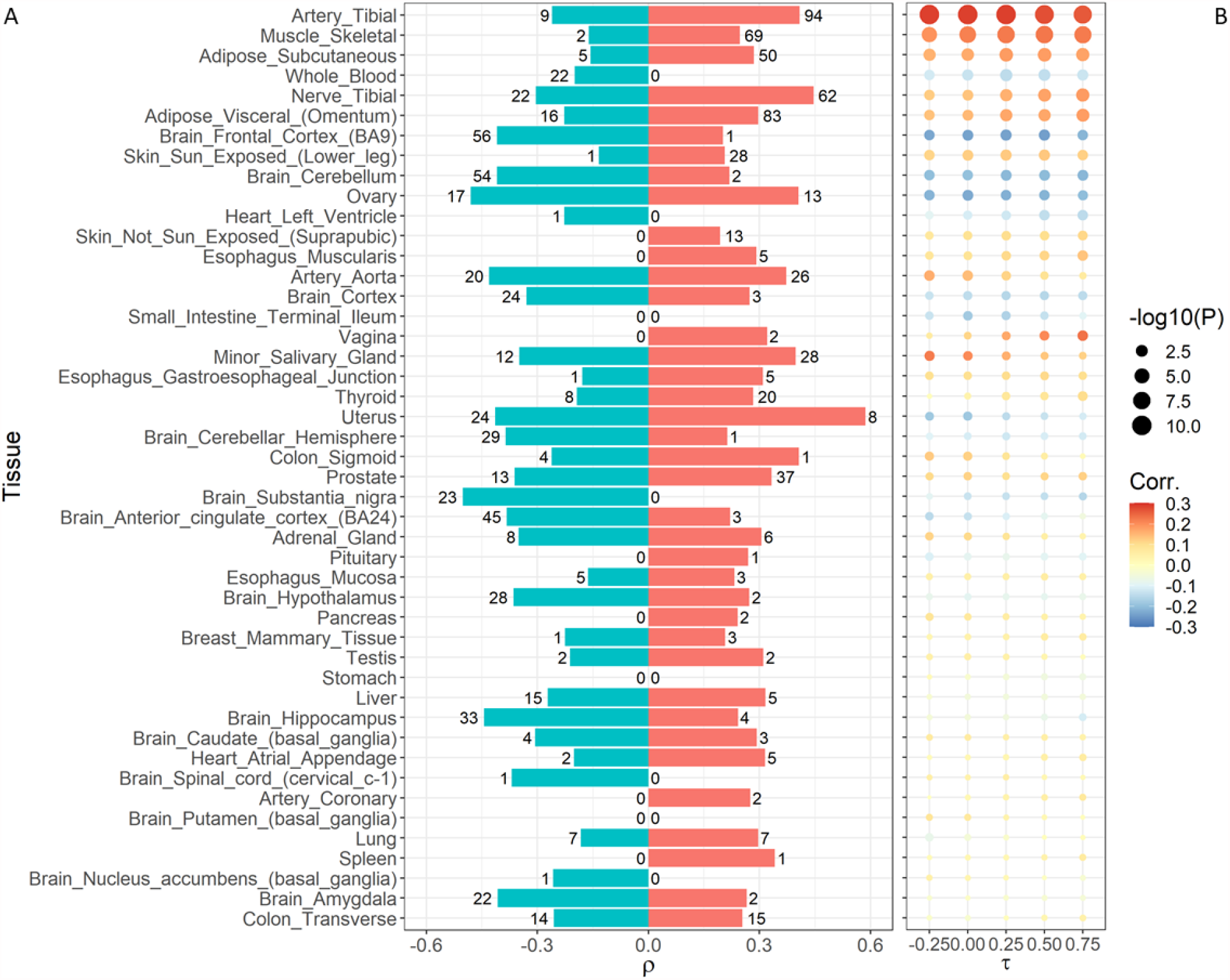
The number of receptors significantly correlated with age in the GTEx data. (A) The range of significant individual receptor/age correlation ρ is shown for each tissue. Numbers next to the bars denote the number of receptors that are significantly positively (red bars) or negatively (green bars) associated with age, respectively. Missing bars indicate the absence of a significant correlation. (B) The age dependency of gene expression between tissues and composite receptor score (CRS) based on the genes coding for cell surface proteins (rows) are shown. Tissues are ranked based on correlation significance with parameter τ = 0.25. Colors refer to the positive (red) and the negative (blue) correlation between age and CRS. The size denotes the FDR in – log_10_ (adj. P-Value).

Among the STSPRs, ectodysplasmin A2 receptor (*EDA2R*) is significantly correlated with age in 39 of the 46 tissues in GTEx analyzed here. Its gene expression level is consistently increased with age across all these tissues (**Table S5**). *EDA2R* is a TNF receptor family member associated with the Nuclear Factor Kappa B (NF-κB) and p53 signaling pathways^36^. *EDA2R* has been identified as a strong candidate gene for lung aging in the context of COPD with additional age association in adipose tissue, artery, heart, muscle, and skin tissue^37^. It is also a target of ACE2 overexpression. Among the STSPRs, *EDA2R* shows the most significant positive correlation in 24 tissues in GTEx, including the tibial artery, subcutaneous fat, tibial nerve, adipose visceral (omentum), or the frontal cortex. EDA2R is a member of the M4-M26 module branch and key regulator in the daughter module M1602.

Other age-associated receptors are *SLC22A15, PSEN1, CD69*, and *ENTPD3. SLC22A15* is positively and negatively correlated with age in 13 and 6 tissues, respectively. It is a member of the prototypical carnitine and ergothioneine transporters^38^ and is associated with many complex lipids that are not characteristic of any other SLC22 transporter^39^. *SLC22A15* facilitates tumorigenesis in colorectal cancer cells. Overexpression of *SLC22A15* leads to an increase in cell proliferation and cell colony formation capacity^40^. Presenilin 1 (*PSEN1*) mutations have been linked to an inherited form of Alzheimer’s disease. *PSEN1* is negatively correlated with age in 16 tissues but positively correlated with age in two brain tissues, the amygdala, and the hippocampus. Presenilins potentially regulate amyloid precursor protein (*APP*) by modulating gamma-secretase, an enzyme that cleaves *APP*. It is further known that the presenilins function in the cleavage of the Notch receptor. *CD69*, a member of the calcium-dependent lectin superfamily of type II transmembrane receptors, is only positively correlated with age in 17 tissues. *CD69* is an early activation marker expressed in hematopoietic stem cells, T cells, and many other cell types in the immune system. Expression of the encoded protein is induced upon activation of T lymphocytes and may play a role in proliferation. Furthermore, the protein may act to transmit signals in natural killer cells and platelets. *CD69* mRNA expression is only positively correlated with age. Thus, we would expect an increased expression with age. However, *CD69* expression and its age-dependency are controversial. CD4+ and CD8+ lymphocytes derived from elderly persons had reduced CD69 surface expression compared to young persons^41^. On the other hand, CD69 enhances the immunosuppressive function of regulatory T-cells in an IL-10 dependent manner^42^. This behavior would fit the hypothesis of a compromised immune response in the elderly. Ectonucleoside triphosphate diphosphohydrolase 3 (*ENTPD3*) is another gene with age-dependent expression. Its expression is positively correlated with age in the tibial artery and skeletal muscle and negatively correlated in 14 tissues. *ENTPD3* encodes a plasma membrane-bound divalent cation-dependent E-type nucleotidase. The encoded protein is involved in regulating extracellular levels of ATP by its hydrolysis (to ADP) and other nucleotides. *ENTPD3* is a key regulator in the M2-M9-M66 branch of modules. Number 4 ranked module M66 (**Figure 3D**) is enriched for macrophages/neutrophils (ARCHS^4^ FET P-value = 0.014, 1.6 FE).

### Age dependency of a systemic SARS-CoV-2 response

Network neighborhoods of several STSPRs such as *ENTPD3, GABRP*, and *EPHA6* are enriched for the SARS-CoV-2 induced DEG signatures from human patient lung samples. The *GABRP* mRNA level is positively correlated with age in three tissues (subcutaneous fat, lung, minor salivary gland) and negatively correlated with age in three other tissues (tibial nerve, not sun-exposed skin, small intestine terminal ileum). *EPHA6*, a member of the M2-M9 branch (**Figure 1A** and **Figure 3C**), promotes angiogenesis^43^ and regulates neuronal and spine morphology^44^. The network neighborhood of *EPHA6* is enriched for pentose and glucuronate interconversion, glucuronidation, and systemic lupus erythematosus (FET P-values < 7.5e-03). *EPHA6* mRNA level increases with age in six tissues (artery aorta, cerebellar brain hemisphere, brain cerebellum, esophagus gastroesophageal junction, esophagus mucosa, and ovary). It decreases in four tissues (brain amygdala, brain cortex, brain hippocampus, and brain hypothalamus). Interestingly, *ACE2* mRNA level increases with age in five tissues (adrenal gland, lung, ovary, stomach, and uterus tissue) and decreases in three tissues (aorta artery, minor salivary gland, and tibial nerve) (**Table S3**).

We further investigated the potential age dependencies of STSPRs in biological processes realized by MEGENA co-expression modules. For this purpose, we have identified network modules enriched for tissue-specific age-correlated STSPR. The 3,227 strong generic transcription module M4 is enriched for both positive and negative correlated STSPRs. M4 is enriched for positive age-correlated STSPR in prostate (FET P-value = 0.015, 1.85 FE) and for negative age-correlated STSPR in liver (FET P-value = 0.0015, 2.77 FE). We have identified the M4-M27 branch with signaling functions underlying COVID-19 (**Figure 1A** shows the M4-M27-M276 branch). Using blood cell type signatures, we found that M4 is enriched for neutrophils (FET P-value = 0.037, 3.0 FE). Neutrophil-mediated innate immune responses against pathogens in the lungs determine the outcome of infection; insufficient neutrophil recruitment can lead to life-threatening infection, although an extreme accumulation of neutrophils can result in excessive lung injury associated with inflammation^45^. Such a massive intra-alveolar neutrophilic infiltration has been observed in COVID-19 patients with a longer clinical course, likely due to superimposed bacterial pneumonia^46^.

Other enriched modules involve number 66 ranked M26 (positive age-correlated STSPRs in adrenal gland: FET P-value = 1.32e-04, 6.88 FE), and number 35 ranked M10 (negative age-correlated STSPRs in mammary breast tissue: FET P-value = 0.069, 6.10 FE). M26 is another child of M4 with cell cycle (M/G1 transition) function.

We also analyzed the dependence of the STSPRs on age in each tissue in the GTEx by computing correlations between differential expression of the STSPRs in COVID-19 and correlations between the STSPRs and age in each tissue in the GTEx (termed STSPR differential expression and age dependence (STSPR-DEAD) score; see details in **Methods and Table S6**). The subcutaneous fat, tibial artery, the substantia nigra, esophagus gastroesophageal junction, and liver show the strongest STSPR-DEAD score. A heatmap of STSPR-DEAD scores between 46 tissues and 7 sample types is shown in **Figure 5A**. Many tissues have negative STSPR-DEAD scores. Examples are tibial artery (ρ = 0.32, p=0.029; **Figure 5B**), liver (ρ = 0.38, p=4.4e-05; **Figure 5C**) and esophagus gastroesophageal junction (ρ = -0.39, p=1.4e-03; **Figure 5D**). The substantia nigra has the strongest positive STSPR-DEAD score and possesses the highest correlation coefficient in absolute terms with DEGs (DEGs from combined BALF and lung biopsies, ρ = -0.32, not shown).

**Figure 5.**
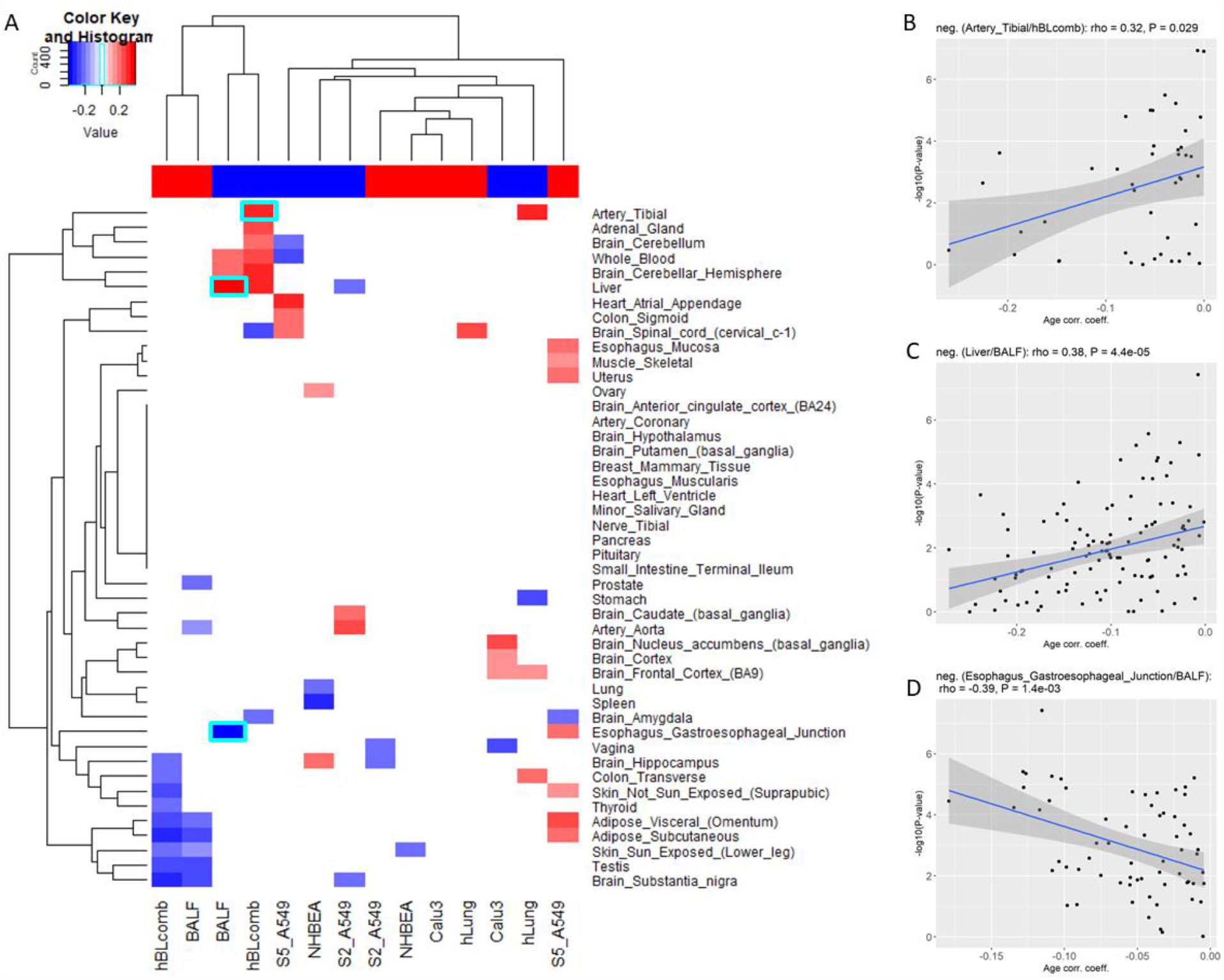
Correlation between the surface receptors’ differential expression in SARS-Cov-2 infection and their tissue-specific age dependence. (A) A heatmap of correlation coefficients after tissue age effect (STSPR-DEAD, see text) and DEGs correlation is shown. Only the correlation coefficients with nominal P ≤ 0.05 are shown. The top color bar indicates the direction of the STSPR-DEADs, with red denoting positive STSPR-DEADs and blue referring to negative STSPR-DEADs. Tiles with cyan boundary indicate select tissue/DEG pairs. (B-D) Dot plots between STSPR-DEAD and DEGs of select tissues with best correlation coefficients are shown: (B) Artery tibia against combined BALF/lung biopsy DEGs, (C) Liver against BALF, and (D) Esophagus Gastroesophageal Junction against BALF.

We have further validated the dependence of the STSPRs on age in GTEx tissues with data from the NYU COVID-19 cohort. Figure S4A shows the heatmap between 46 tissues, 3 sample types, and one combined data set (Xsq), corresponding to Figure 5A. Figure S4B depicts a plot between STSPR-DEAD and DEGs of esophagus gastroesophageal junction against HL (ρ = -0.49, p=3.2e-05) corresponding to Figure 5D.

To explore the gene expression changes of STSPRs with age, we have separated GTEx donors into two cohorts: a young (≤ 45yrs) cohort and an old cohort (≥ 60yrs). Gene expression was then adjusted to compare the difference between these two cohorts (see methods). In subcutaneous fat and tibial artery, the young cohort showed a lower gene expression level, while a higher level of gene expression in the elder cohort. This pattern can also be seen in the esophagus gastroesophageal junction, skeletal muscle (**Figure S5**).

Overall, we found a clear age-effect of genes coding for cell surface proteins and receptors that are potentially utilized by SARS-CoV-2. In particular, we have identified that STSPRs showed stronger age-dependency in the tibial artery, skeletal muscle, adipose, and brain tissues. Such an age-dependent effect could potentially contribute to the elevated severity of COVID-19 in the elderly.

## Discussion

In the present study, we focus on the biological processes and key regulators modulating the host response to SARS-CoV-2 infections. Our multiscale network analysis of the gene expression data from both patient samples and cell lines has revealed network structures and key regulators underlying the host response to SARS-CoV-2 infection.

Essential aspects in the COVID-19 pathology are the biological processes hijacked by the virus for its advantage. Expression of the *ACE2* receptor on the host cell and binding of the viral Spike protein for cell entry are among the first steps. Other processes beneficial for the virus may be staged by *ACE2* expression and triggered by the binding process. *CD300e* and its interacting partner *TYROBP* trigger pro-inflammatory cytokines and prevent apoptosis, an essential process controlled by many viruses. On the other hand, severe inflammation significantly contributes to the pathology of COVID-19 disease. Other potential surface protein host-factors are *CD81* and *EGFR*. Additional surface proteins are *CEACAM1* and *LAMP3*. Multiple cellular activities have been attributed to *CEACAM1*, including differentiation and arrangement of three-dimensional tissue structure, angiogenesis, apoptosis, tumor suppression, metastasis, and the modulation of innate and adaptive immune responses. *LAMP3*, however, plays a critical role in the autolysosome fusion process. It activates the PI3K/AKT pathway, which is necessary for SARS-CoV to establish infection.

We have further investigated the age-dependence of receptors’ expression as clinicians have observed a severe disparity in survival between old and young COVID-19 patients. We have identified a strong correlation between tissue age-dependency and SARS-CoV-2 infection-induced receptor expression in subcutaneous fat, tibial artery, brain substantia nigra, esophagus gastroesophageal junction, and liver. However, the exact contribution of specific receptors’ age-dependency on the disease’s pathology requires additional investigation. We have also identified specific genes potentially related to age-specific expression and response in SARS-CoV-2 infections. *EDA2R* expression is significantly positively correlated with age in 24 of 46 tissues in GTEx, including the tibial artery, subcutaneous fat, tibial nerve, adipose visceral (omentum), the frontal cortex, or lung. Concerning lung, *EDA2R* has been associated with aging in the context of the chronic inflammatory disease COPD^37^. This particular gene is another target of ACE2 overexpression, potentially affected as a response to SARS-CoV-2 infection. Other targets with age-dependent expression are *CD69, ENTPD3, EPHA6, GABRP, PSEN1*, and *SLC22A15*. Noteworthy are *CD69* and *ENTPD3*. As a surface receptor on immune cells and involved in signal transduction, *CD69* is an integral component of immune system functions. With its positive correlated, age-dependent expression and its known modulation of immunosuppressive function of regulatory T-cells^42^, *CD69* may contribute to compromising immune response in the elderly during SARS-CoV-2 infection. The ectonucleotidase *ENTPD3* shows a protective role in intestinal inflammation^47^ and maybe another factor of the age-dependent immune reaction during COVID-19. It is a key regulator with a network neighborhood enriched for genes responding to ACE2 overexpression.

In conclusion, our analyses presented here suggest that SARS-CoV-2 utilizes multiple novel receptors for entry and spawns a unique response in the host system. Novel hypotheses involving the utilization of cell surface receptors and their age-dependent expression offer new insights into the molecular mechanisms of SARS-CoV-2 infection and pave the way for developing new therapeutic intervention against COVID-19.

## Methods

### RNAseq Analysis

Raw reads were obtained from the Beijing Institute of Genomics (BIG) Data Center (https://bigd.big.ac.cn/) under the accession number CRA002390. BALF RNAseq data from healthy subjects were obtained from NCBI/SRA (SIB028/SRR10571732, SIB030/ SRR10571730, and SIB036/ SRR10571724). The RNAseq data were aligned to the Homo sapiens reference genome GRCh38/hg19 using the Star aligner v2.7.0f with modified ENCODE options, according to Xiong *et al*..^15^ Raw read counts were calculated using featureCounts v2.0.1. Raw read counts after Star alignment and featureCounts, as well as obtained from GSE147507, were normalized using edgeR/voom (v3.32.1 with R v4.0.0).

### Identification of differentially expressed genes

We used the negative binomial models together with the empirical Bayes approach as implemented in the edgeR-package^48^ to identify differentially expressed genes (DEGs). We considered an absolute fold change of 1.5 or higher and an FDR ≤ 0.05 as significant throughout the paper.

### Gene co-expression network analysis

Multiscale Embedded Gene Co-Expression Network Analysis (MEGENA) ^20^ was performed to identify host modules of highly co-expressed genes in SARS-CoV-2 infection. The MEGENA workflow comprises four major steps: 1) Fast Planar Filtered Network construction (FPFNC), 2) Multiscale Clustering Analysis (MCA), 3) Multiscale Hub Analysis (MHA), 4) and Cluster-Trait Association Analysis (CTA). The total relevance of each module to SARS-CoV-2 infection was calculated by using the Product of Rank method with the combined enrichment of the differentially expressed gene (DEG) signatures as implemented: *G*_*j*_ = П_*i*_ *g*_*ji*_, where, *g*_*ji*_ is the relevance of a consensus ***j*** to a signature ***i;*** and *g*_*ji*_ is defined as *(max*_*j*_ *(r*_*ji*_) + 1 - *r*_*ji*_)/∑_*j*_ *r*_*ji*_, where *r*_*ji*_ is the ranking order of the significance level of the overlap between the module ***j*** and the signature.

### Identification of enriched pathways and key regulators in the host modules

To functionally annotate gene signatures and gene modules identified in this study, we performed an enrichment analysis of the established pathways and signatures–including the gene ontology (GO) categories and MSigDB–and the subject area-specific gene sets–including, Inflammasome, Interferome, and InnateDB. The hub genes in each subnetwork were identified using the adopted Fisher’s inverse Chi-square approach in MEGENA; Bonferroni-corrected p-values smaller than 0.05 were set as the threshold to identify significant hubs.

### Network enrichment

Fisher’s Exact Test (FET) was performed to determine the overlap between network neighborhoods of potential key regulators (target) and an input DEG signature. For each target in the network in the 95 percentile of node strength after MEGENA, the genes in the network neighborhoods between one and four steps away from the target were intersected with the DEG signature. MEGENA networks were tested with DEGs of all systems for further analysis (see **the main text**). Cumulative network enrichment scores *s* = 1/*n* · ∑_*i*_ - log_10_ *P*_*i*_ based on individual FET P-values for each target were calculated. *n* is the number of realizations (i.e., the number of different neighborhoods and systems used to calculate the particular score).

### GTEx data preprocessing

We downloaded GTEx v8 data^49^ from the Database of Genotypes and Phenotypes (dbGaP) under accession phs000424.v8.p2. For all the available tissues, we selected those with at least 80 samples and samples with more than 20 million mapped reads and greater than a 40% mapping rate. Cell line data were removed from our analysis. Only genes with expression > 0.1 Transcripts Per Million (TPM) and aligned read count of 5 or more in more than 80% samples within each tissue were used for aging gene identification. Expression measurements for each gene in each tissue were subsequently inverse-quantile normalized to the standard normal distribution to reduce the potential impact of outlier gene expression values. Our final dataset included samples from 46 tissue types. The sample size for each tissue ranged from 114 to 706, with an average of 315 samples.

### Linear regression model for age and sex-associated gene detection

We implemented a linear regression model to identify age-associated gene expression (Eq. 1) ^50^.

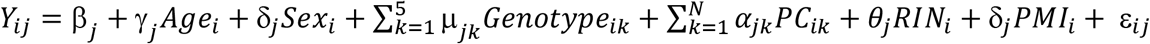(Eq. 1). In this model, *Y*_*ij*_ is the expression level of gene j in sample i, *Age*_*i*_ denotes the donor age of sample i, *Sex*_*i*_ denotes the donor sex for sample i, *Genotype*_*ik*_ (*k* ∈ (1,2,3,4,5)) denotes the value of the k-th principal component value of the genotype profile for the i-th sample, *PC*_*ik*_ (*k* ∈ (1, …, *N*) denotes the value of the k-th principal component value of gene expression profile for the i-th sample, N is the total number of top PCs under consideration, *RIN*_*i*_ denotes the RIN score of sample i, *PMI*_*i*_ denotes the PMI of sample i, *ε*_*ij*_ is the error term, *γ*_*j*_, *δ*_*j*_, *μ*_*jk*_, *α*_*jk*_, *θ*_*j*_, *δ*_*j*_ are the regression coefficients for each covariate. The corresponding correlation coefficients and p-values (adjusted with BH ^51^ method) were then calculated for all genes; FDR values < 0.05 were considered as significant age-associated genes. Several covariates (such as genotype PCs and PEER factors) we adjusted in the regression model were selected following the method used by GTEx consortium ^49^. From the consortium’s analysis, the top five genotype PCs were considered sufficient to capture the major population structure in the GTEx dataset and were used for the consortium paper.

### Adjust gene expression for age analysis

We used a linear regression model to adjust gene expression (Eq. 2).*Y*_*ij*_ = b_*j*_ + *δ*_*j*_ *Sex*_*i*_ + *μ*_*j*_ *Platform*_*i*_ + *θ*_*j*_ *RIN*_*i*_ + *δ*_*j*_ *PMI*_*i*_ + *ε*_*ij*_ (Eq. 2). We regressed out the following confounding factors to obtain adjusted gene expression, which include *Sex*_*i*_ : the sex of donor for sample i, *Platform*_*i*_ : the value of the platform for the i-th sample, *RIN*_*i*_ : the RIN score of sample i, and *PMI*_*i*_ : the PMI of sample i.

Expression measurements for each gene in each tissue were inverse-quantile normalized to follow the standard normal distribution to reduce the potential impact of outlier gene expression values. Composite receptor score (CRS) was then calculated for each receptor in each sample (Eq.3). *CRS*(*Y*_*i*_) = *sum*{*sign*(*X*_*ij*_, *τ*)} where 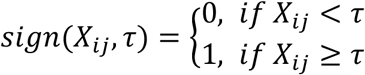(Eq. 3). In this equation, *CRS(Y*_*i*_ ) is the composite score of sample i, *X*_*ij*_ is the expression level of gene j in sample i, *τ* is the test score. We have tested *τ* with -0.25, 0, 0.25, 0.5, 0.75, and 1, spearman correlation coefficients and p-values (adjusted with BH method) were subsequently calculated between CRS score and age. *τ* = 0.25 showed the overall best correlation and p-value between CRS and age (**Table S6**). We termed this correlation coefficient between SARS-CoV-2 surface protein receptors (STSPRs) CRS and age, STSPR differential expression, and age dependence (STSPR-DEAD) score.

## Acknowledgments

This work was funded by grants to CVF from the National Institutes of Health (R21 AI149013).

The Genotype-Tissue Expression (GTEx) Project was supported by the [Common Fund](https://commonfund.nih.gov/GTEx) of the Office of the Director of the National Institutes of Health and by NCI, NHGRI, NHLBI, NIDA, NIMH, and NINDS.

## Author Contributions

CVF and BZ conceived and designed the study. CVF, LZ, QW, ZX, and SV analyzed the data. ZT provided insights into age dependency. CVF and BZ wrote the paper.

## Competing Interests

The authors declare no competing interests.

## Data Availability

The datasets analyzed during the current study are available from the corresponding author on reasonable request.

## GLOSSARY

BALF: Bronchoalveolar lavage fluid
CoV: Coronavirus
COVID-19: Coronavirus disease 2019
CRS: Composite receptor score
DEG: Differentially expressed gene
FC: Fold change
FDR: False discovery rate
FET: Fisher’s exact test
IFN: Interferon
ISG: Interferon stimulated gene
NHBE: Normal human bronchial epithelial (cells)
PRR: Pattern recognition receptor
STSPR: SARS-CoV-2 triggered surface protein receptor
STSPR-DEAD: STSPR differential expression and age dependence (score)

## Notes

### Competing Interest Statement

The authors have declared no competing interest.

