## Supplemental Material for "Tissue Specific Age Dependence of the Cell Receptors Involved in the SARS-CoV-2 Infection"

Christian V. Forst[1,2,3,4], Lu Zeng[1], Qian Wang[1,2,3], Xianxiao Zhou[1,2,3], Sezen Vatansever[1,2,3], Zhidong Tu[1,3], Bin Zhang[1,2,3,5,*]

[1]Department of Genetics and Genomic Sciences, Icahn School of Medicine at Mount Sinai, 1425 Madison Avenue, NY 10029
[2]Mount Sinai Center for Transformative Disease Modeling, Icahn School of Medicine at Mount Sinai, 1470 Madison Avenue, NY10029
[3]Icahn Institute for Data Science and Genomic Technology, Icahn School of Medicine at Mount Sinai, 1425 Madison Avenue, NY10029-6501, USA.
[4]Department of Microbiology, Icahn School of Medicine at Mount Sinai, One Gustave L. Levy Place, New York, NY 10029
[5]Department of Pharmacological Sciences, Icahn School of Medicine at Mount Sinai, 1425 Madison Avenue, NY 10029

*Corresponding author:

Bin Zhang, PhD

Professor, Department of Genetics & Genomic Sciences, Icahn Institute of Genomics and Multiscale Biology

Icahn School of Medicine at Mount Sinai

1470 Madison Avenue, Room 8-111

New York, NY 10029

(O) 212-824-8947

***Network enrichment***.

We tested the network neighborhood of each gene in the network for enrichment by previously identified DEG signatures (e.g., SARS-CoV-2 DEGs obtained from patients and cell data), termed a network enrichment score (NWes) based on the cumulative P-values. The network neighborhood of a given gene $x$ is comprised of the genes that are within N-layers (N = 1, 2, …) away from $x$ in a given network. We then calculated a compound network enrichment score (NWes) based on the cumulative P-values after enrichment using data from all samples. We then rank-ordered these key regulators based on their NWes.

For further functional assessment of the MEGENA modules, we used a set of gene signatures, such as transcription factors and their binding sites, miRNA targets, ISGs, inflammasome, and cell- and tissue type. We were also employing the *ACE2* overexpression (ACE2oe) data from Blanco-Melo *et al*., 2020[1] to identify processes affected by *ACE2* expression. ACE2oe expression signatures were called by a hierarchical linear model (hLM) using Limma[2] identified genes with a different expression pattern between infected vs. control wildtype cells and ACE2oe infected vs. control cells. We have identified 2792 unique genes with this expression pattern with FDR ≤ 0.05 and a cutoff of 1.5 of the absolute value of the fold change (FC), and 1606 genes using an absolute FC of 2 or more. For a definition of FC in the hLM case, see Methods.

**Key molecular regulators during SARS-CoV-2 infection**

Other top-ranked key regulators not reported in the main text are roundabout guidance receptor 3 (*ROBO3*), sperm-associated antigen 16 (*SPAG16*), and transmembrane p24 trafficking protein 9 (*TMED9*). *ROBO3* is a member of the immunoglobulin transmembrane receptor superfamily. The *ROBO3* gene regulates axonal navigation at the ventral midline of the neural tube. It is down-regulated in lung samples (BALF: 35.7 fold, FDR = 2.1e-04; lung biopsy:

10.5 fold, FDR = 2.6e-06) and slightly up-regulated in A549 cells (1.8 fold, FDR = 0.019). *SPAG16* is upregulated in lung samples (BALF: 12.8 fold, FDR = 2.6e-04; lung biopsy: 6.8 fold, FDR = 8.5e-8) and down-regulated in Calu3 cells (1.8 fold, FDR = 0.012). SPAG16 potentially has immune system functions and may play a role in auto-immune diseases. It is a target of the humoral autoimmune response in multiple sclerosis[3]. It also influences matrix metalloproteinase (MMP-3) regulation and protects against joint destruction in autoantibody-positive rheumatoid arthritis[4]. *TMED9* is a member of a family of genes encoding transport proteins located in the endoplasmic reticulum (ER) and the Golgi. It is down-regulated in both lung samples and cell lines (BALF: 6.8 fold, FDR = 5.9e-04; lung biopsy: 2.8 fold, FDR = 7.3e-08; Calu3: 1.8 fold, FDR = 5.4e-03; A549: 1.4 fold, FDR = 8.3e-03). TMED9 is an ER cargo receptor that regulates intracellular vesicle biogenesis. It may also be involved in pathological Golgi-lysosome transport[5]. The reduction of such a cargo receptor may disrupt the Golgi system and induce retrograde transport from the cell surface back to the ER[6]. Down-regulation of TMED9 and the evidence of lysosome utilization by SARS and MERS viruses[7] may support this hypothesis.

**Receptors and host-factors required for viral entry**

The best-ranked ACE2oe network enriched targets are *CLOCK*, *CD300E*, *CD81*, *C14orf119*, and *CTSZ*.  Clock circadian regulator (*CLOCK*) plays a central role in the regulation of circadian rhythms. *CLOCK* , a transcription factor, is up-regulated in BALF and A549 samples (BALF: 2.6 fold, FDR = 0.011; A549: 1.3 fold, FDR = 0.05). Thus *CLOCK* does not function as a surface protein. *CD300e* is a member of the *CD300* glycoprotein family of transmembrane cell surface proteins expressed on myeloid cells. It is up-regulated in lung samples (BALF: 18.5 fold, FDR = 2.7e-03; lung biopsy: 2.7 fold, FDR = 0.030). The protein interacts with the TYRO protein tyrosine kinase binding protein (*TYROBP*) and is thought to act as an activating receptor. Activation via *CD300e* provided survival signals that prevented monocyte and Myeloid dendritic cells apoptosis, triggered the production of pro-inflammatory cytokines, and upregulated the expression of cell surface co-stimulatory molecules in both cell types[8]. The expression and function of human *CD300* receptors on blood circulating mononuclear cells are distinct in neonates and adults[9], potentially contributing to the difference in clinical outcome after COVID-19 infection. Another ACE2oe network enriched target is *CD81*, with down-regulation in lung samples and cell-lines (BALF: 95.5 fold, 1.6e-04; lung biopsy: 4.0 fold, 0.025; A549: 1.3 fold, FDR = 8.2e-03; Calu3: 1.5 fold, FDR = 0.026). The protein encoded by this gene is a member of the transmembrane 4 superfamily, also known as the tetraspanin family.

Most of these members are cell-surface proteins that are characterized by the presence of four hydrophobic domains. The proteins mediate signal transduction events that play a role in regulating cell development, activation, growth, and motility. This encoded protein is a cell surface glycoprotein that is known to complex with integrins. Furthermore, *CD81* is an entry co-receptor for the Hepatitis C virus. *CD81* is the only ACE2oe target which network neighborhood is significantly enriched for SARS-CoV-2 signatures, yielding a rank #79 based on NWes. Thus, CD81 is potentially a novel host cell receptor that SARS-CoV-2 requires for entry and, thus, a therapeutic target. *C14orf119* is another ACE2oe enriched target, up-regulated in human lung biopsy samples (1.4 fold, FDR = 0.049), with unknown function.

Cathepsin Z (*CTSZ*) is a lysosomal cysteine proteinase and member of the peptidase C1 family. It is down-regulated in lung samples (BALF: 18.6 fold, FDR = 0.011; lung biopsy: 1.9 fold, FDR = 2.4e-03) and slightly up-regulated in A549 (1.2 fold, FDR = 0.040). Singh et al., 2020 hypothesized that cathepsins are among other factors that facilitate SARS-CoV-2 entry into the host cell[10]. The epidermal growth receptor *EGFR* is a transmembrane glycoprotein and present on the cell surface of epithelial cells. It is significantly upregulated in lung samples (BALF: 74.9 fold, FDR = 6.82e-04 and lung biopsy: 2.9 fold, FDR = 0.022), A549 (2.3 fold, FDR = 2.27e-03) and Calu3 (2.9 fold, FDR = 0.011) cells. EGFR is a host-factor for hepatitis C virus entry[11]. Respiratory viruses induce EGFR activation, suppressing IFN regulatory factor (IRF) 1–induced IFN-λ, and antiviral defense in airway epithelium[12]. Thus, *EGFR* may not be required for SARS-CoV-2 entry, but it may be a potential host factor for the viral life cycle.

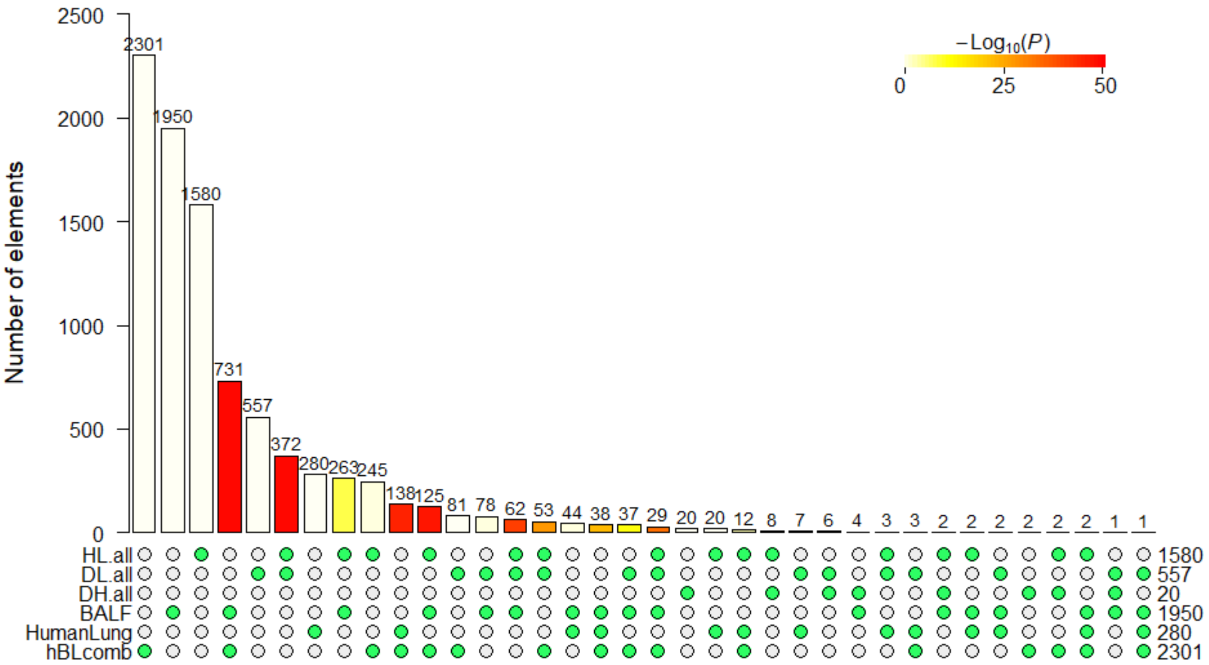


**Figure S1: Super Exact Test between different patient-derived DEGs**. Multi-set intersections between DEGs identified in this publication (BALF, HumanLung, hBLcomb) and derived from the NYU COVID-19 cohort (DH, DL, HL) are shown. DH, DL, and HL denote comparisons between different clinical COVID-19 outcomes: DH…death vs. prolonged mechanical ventilation (>28 days), DL…death vs. short mechanical ventilation (<28 days), HL prolonged vs. short mechanical ventilation. Vertically arranged green circles indicate corresponding intersections (or lack thereof).

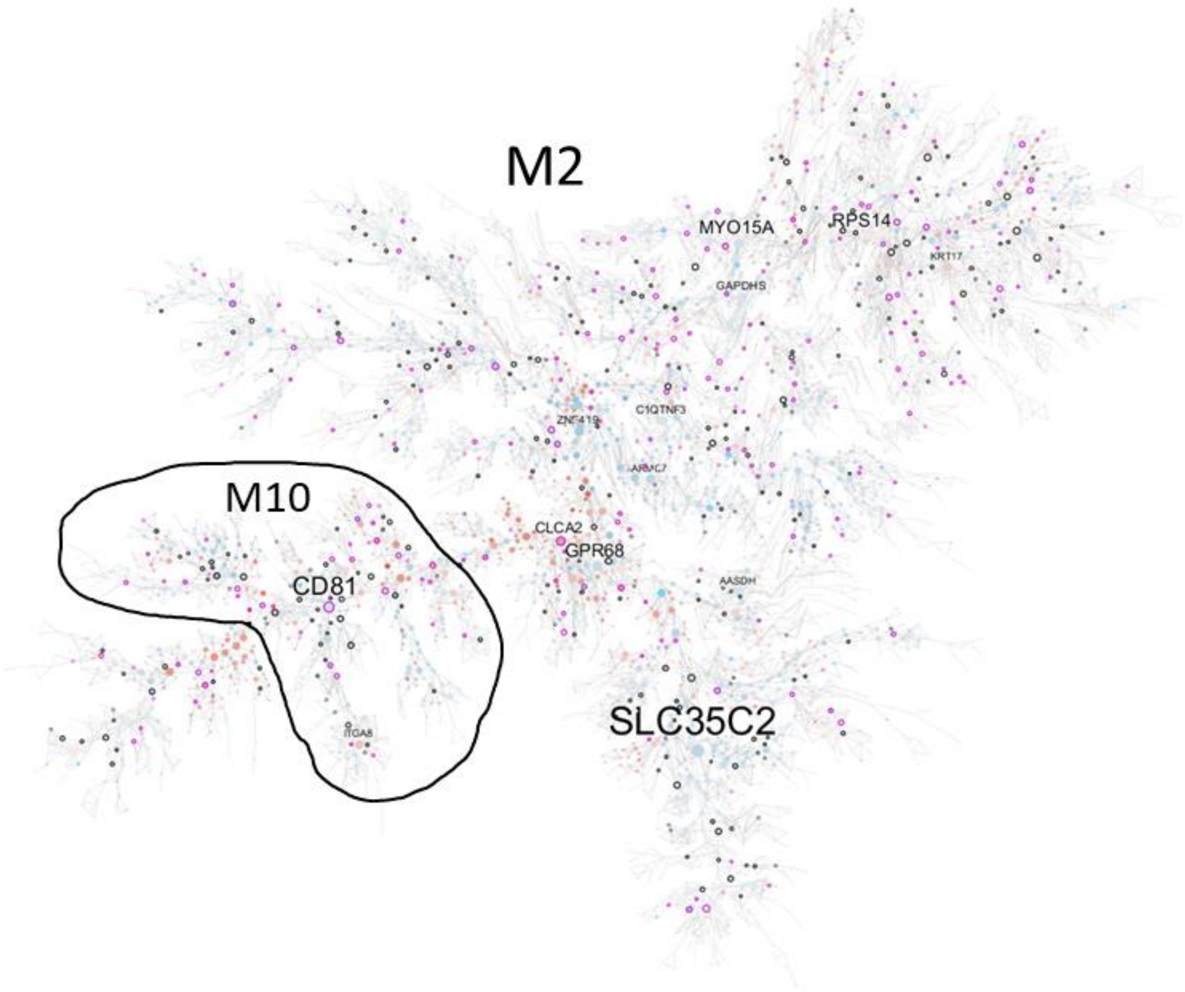


**Figure S2: Gene co-expression modules associated with SARS-CoV-2 infection**. M2 (#1), the parent module of M10 (highlighted), is shown. The node color indicates a directional response. Red nodes are upregulated, blue nodes are downregulated after infection. Diamond-shaped nodes indicate key regulators. The nodes with a black border denote genes significantly responding to ACE2 overexpression with fold change (FC) of 1.5 or higher. Purple borders indicate ACE2oe responding genes with FC ≥ 2.

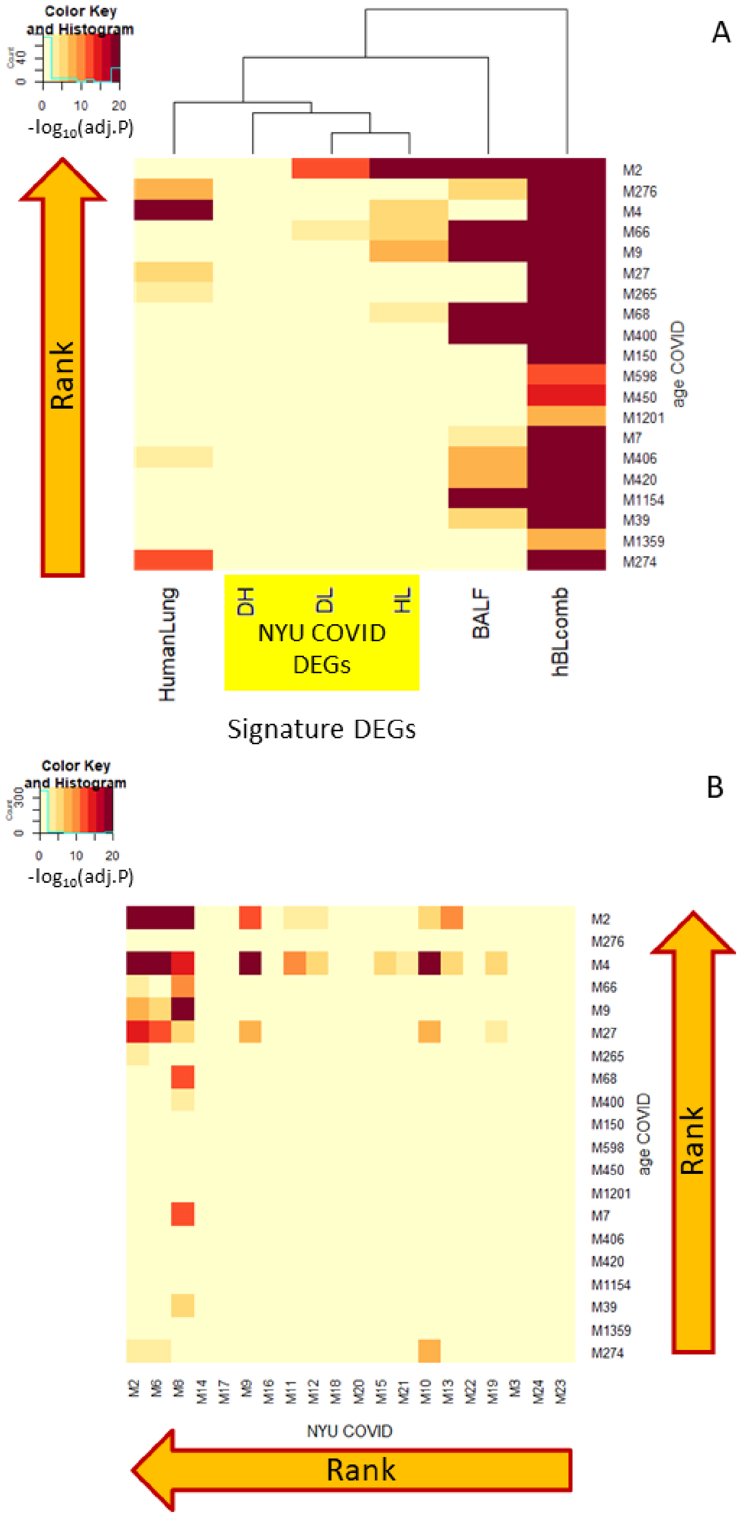


**Figure S3: Module-module overlap and enrichment**. (A) The enrichment of 20 best-ranked modules by signature DEGs is shown (with top-ranked module M2). The yellow box indicates DEGs obtained from the NYU COVID-19 cohort). (B) THE FET overlap between modules from this paper and modules derived from the NYU COVID-19 cohort is depicted.

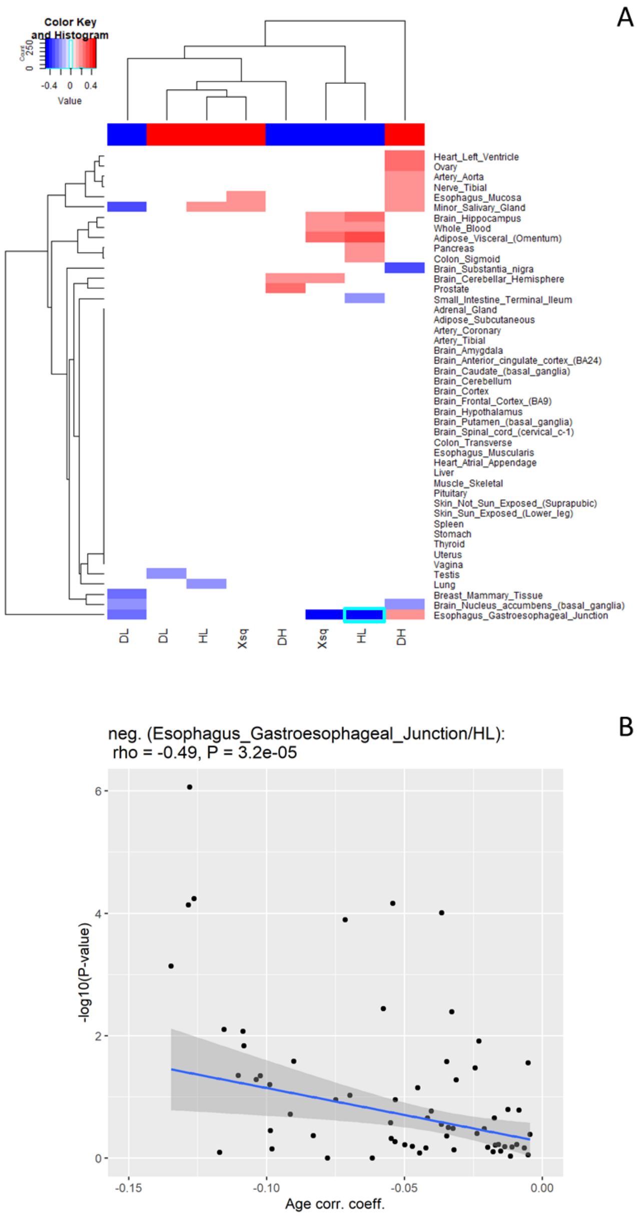


**Figure S4. Validation of the correlation between the surface receptors' differential expression in SARS-Cov-2 infection and their tissue-specific age dependence using data from the NYU COVID-19 cohort.** (A) A heatmap of correlation coefficients after tissue age effect (STSPR-DEAD, see text) and DEGs correlation using NYU COVID-19 patient data is shown. Only the correlation coefficients with nominal P ≤ 0.05 are shown. The top color bar indicates the direction of the STSPR-DEADs, with red denoting positive STSPR-DEADs and blue referring to negative STSPR-DEADs. Rows indicate tissues, columns refer to different comparisons of COVID-19 severity in NYU patients: DH…death vs. high time on a ventilator, DL…death vs. low time on a ventilator, HL… high vs log time on a ventilator. Xsq denotes combined (DH, DL, HL) data after Fisher's combined probability test. The tile with cyan boundary indicates select tissue/DEG pairs. (B) A dot plot between STSPR-DEAD and DEGs of Esophagus Gastroesophageal Junction against HL is shown

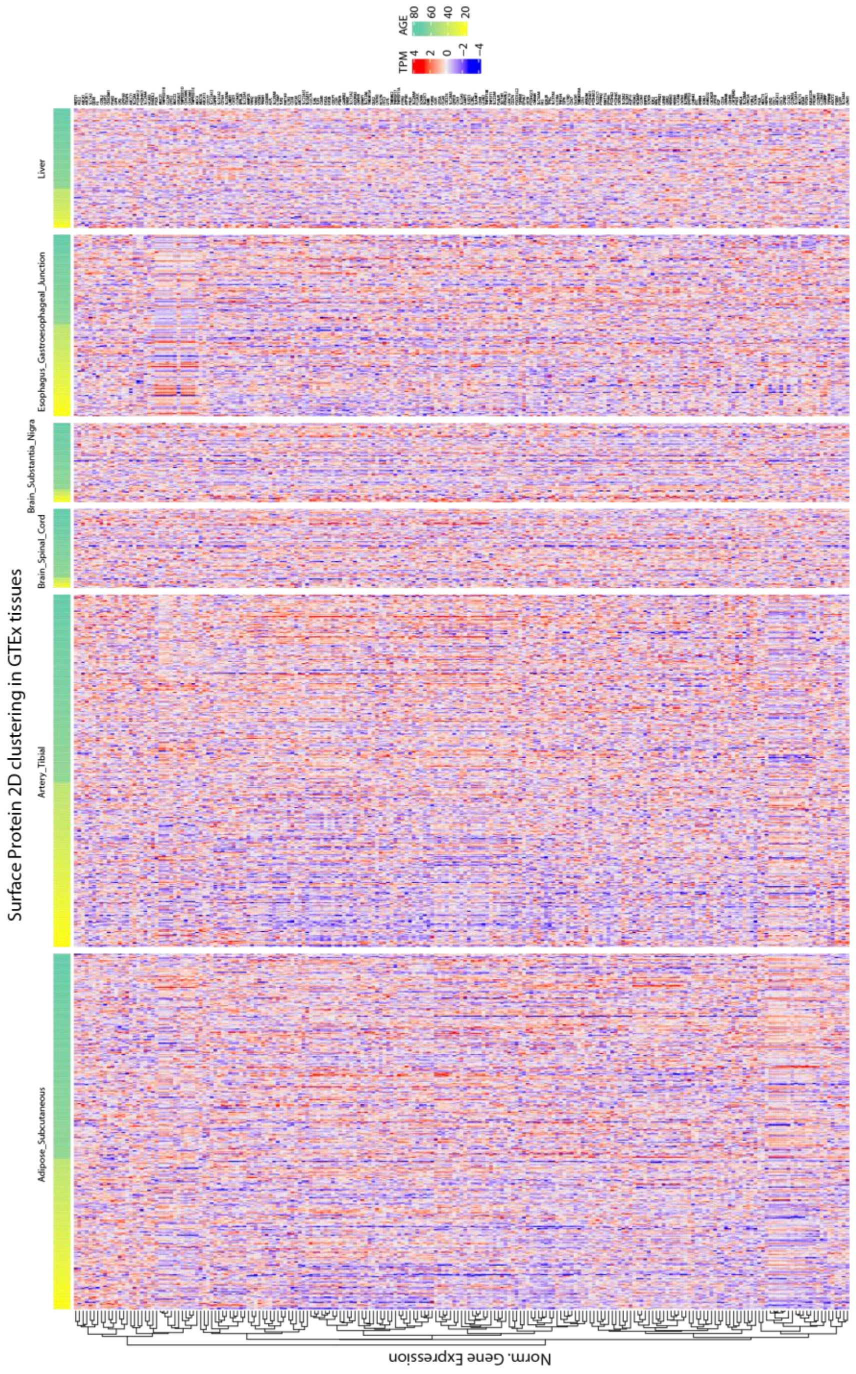


**Figure S5: The age-dependency of cell surface proteins.** The divided heatmap shows the age-dependencies of cell surface proteins in six different tissues (subcutaneous fat, tibial artery, spinal cord, substantia nigra in the brain, esophagus gastroesophageal junction, liver). The yellow-to-green horizontal bars denote age with young (yellow) and old (green). The normalized gene expression in transcripts per million (TPM) is denoted by the red-to-blue color bar with up-regulated (red) and down-regulated (blue) genes.
